# Identifying and Classifying Goals For Scientific Knowledge

**DOI:** 10.1101/2021.05.27.445866

**Authors:** Mayla R. Boguslav, Nourah M. Salem, Elizabeth K. White, Sonia M. Leach, Lawrence E. Hunter

## Abstract

**Motivation:** Science progresses by posing good questions, yet work in biomedical text mining has not focused on them much. We propose a novel idea for biomedical natural language processing: identifying and characterizing the *questions* stated in the biomedical literature. Formally, the task is to identify and characterize **ignorance statements**, statements where scientific knowledge is missing or incomplete. The creation of such technology could have many significant impacts, from the training of PhD students to ranking publications and prioritizing funding based on particular questions of interest. The work presented here is intended as the first step towards these goals.

**Results:** We present a novel ignorance taxonomy driven by the role ignorance statements play in the research, identifying specific goals for future scientific knowledge. Using this taxonomy and reliable annotation guidelines (inter-annotator agreement above 80%), we created a gold standard ignorance corpus of 60 full-text documents from the prenatal nutrition literature with over 10,000 annotations and used it to train classifiers that achieved over 0.80 F1 scores.

**Availability:** Corpus and source code freely available for download at https://github.com/UCDenver-ccp/Ignorance-Question-Work. The source code is implemented in Python.

## 1 Introduction

Science progresses by posing good questions as well as through analysis of experimental results [14, 23, 32]. Until now, biomedical text mining has focused on extracting information from the literature [7] either ignoring [13, 15, 42, 31] or de-emphasizing [44, 20, 27, 21] questions and other statements regarding uncertainty or missing knowledge. The scientific literature is full of statements about knowledge that does not exist, including goals for desired knowledge, statements about uncertainties in interpretation of results, discussions of controversies, and many others; collectively, we term these *ignorance statements*. Ignorance statements are the subject of many passages in nearly every biomedical publication, even when not stated explicitly as a syntactic question [14, 23, 32]. Here we describe a novel natural language processing (NLP) task: identification and characterization of ignorance statements. We present initial results that characterize the extent of ignorance statements in a sample of the full text biomedical literature, provide a theoretically-driven taxonomy of such statements, describe a manually annotated corpus of ignorance statements and their categorization (along with novel annotation guidelines for the task), and demonstrate that it is possible to automatically identify and classify ignorance statements.

An automated approach to identifying and characterizing ignorance statements has a variety of significant use cases. One important potential application is to facilitate interdisciplinary interactions: A collection of formally characterized ignorance statements could be used to identify open questions from other disciplines that new results might bear on, particularly when considering genome-scale data such as transcriptomics. A systematic survey of open scientific questions could be useful to a wide variety of scientists, ranging from graduate students looking for thesis projects to funding agencies tracking emerging research areas. Another potential application is longitudinal analysis, for example, tracking the evolution of research questions over time.

Although this task is novel, it is related to prior NLP efforts to characterize uncertainty, hedging, speculation, and meta-knowledge [13, 15, 42, 31, 44, 20, 27, 21, 6, 9, 2, 3, 8, 39, 33]. It also draws from related work in the philosophy of science regarding how scientists choose problems and approaches, which requires that they characterize their goals for new knowledge [14, 23, 41, 5, 28, 34, 43, 38, 16].

While these ideas and methods are generally applicable across biomedical research, here we focus on one specific area, prenatal nutrition, as a tractable demonstration of feasibility. Our area of focus was selected for its global health significance and a modest-sized but diverse literature. Women are understudied in clinical research [17, 40], especially when pregnant [30, 29], due to ethical and legal considerations and complexities. We hope that even this initial pilot demonstration will be significant, as it has the potential to facilitate new interdisciplinary interactions that could advance the study of this underserved population.

### 1.1 Related Work

Previous work in both NLP and more broadly provides the foundation of our approach, but differs in a variety of important ways. There is no agreed upon name for the phenomena we are calling *ignorance*; prior work has used the related terms *hedging, uncertainty, speculation*, and *meta-knowledge*, each of which has been defined to be somewhat different than our focus here.

Philosophers of science have attended to ignorance statements as a driving force in the selection of research topics and approaches. Thomas Kuhn’s work on the structure of scientific revolutions [23] mentioned the role of unanswerable questions (which he called anomalies) in paradigm shifts. Sylvain Bromberger [5] was the first to argue specifically that questions drive science, and focused on the need for philosophers to address how scientists select topics for research. Biologist Stuart Firestein [14] extended that view, and described an approach to scientific pedagogy based on what he termed *ignorance*, which is where we borrow our term from. Michael Smithson [41] extended Bromberger’s arguments and offered a taxonomy of ignorance from the mathematical and psychological perspectives.

Computational ontologists apply these philosophical principles in a more practical fashion, defining terminologies and relationships intended to describe degrees of evidence or confidence. The Evidence Ontology (ECO) [9], provides a means for representing the types of support for specific assertions. The Confidence Information Ontology (CIO) [3] extends ECO to allow for the specification of varying degrees of confidence based on evidence type. The Scientific Evidence and Provenance Information Ontology (SEPIO) [6] extends that work by providing representational tools to capture the sources of evidence about assertions. The Ontology for Biomedical Investigations (OBI) [2] focuses on representation of planned scientific processes (e.g. experiments) and provides some terms for capturing the goals of those processes, that is the scientific questions being addressed. However, none of these prior ontological efforts is adequate to represent the diverse family of ignorance statements found in the scientific literature.

Natural language processing research has a long history of work on the identification of uncertainty, hedging and speculation, mostly with the goal of excluding such statements from information extraction efforts. Hedging was first defined in linguistics [25] as statements that can be true or false to some extent. Hyland [19] provided a comprehensive linguistic analysis of hedging, specifically focused on the scientific literature. Light et al. [27] conducted a preliminary analysis of speculative language in scientific abstracts to show it is feasible for human readers to annotate. Medlock et al. [31] extends the work of Light et al. by updating the guidelines, annotating full-text articles, and training a classifier. Using the Medlock, et al. corpus, Kilicoglu et al. [20] analyze the linguistic features of speculation, producing a set of lexical cues and syntactic patterns that distinguish speculative language versus not (unhedges). Vincze et al. [42] built a larger corpus, annotating uncertainty and negation as well as identifying the scope (specific text span) that was hedged or negated. Wikipedia has systematically tagged hedged statements as “weasels” [15]. A shared task in 2010 [13] included identifying biomedical hedges in BioScope and the weasels in Wikipedia. Zerva et al. [44] and Kilicoglu et al. [21] build on this work with more elaborate automated approaches to assessing confidence of NLP-based information extraction results.

In contrast to the work on hedging and speculation intended to excise such statements from factual material such as Wikipedia, our goal is to identify ignorance statements in scientific writing in order to analyze and use them directly. Related work includes Patrick et al. [33], who aimed to classify questions in the clinical domain pertaining to electronic patient notes in order to determine if questions were answerable and answer them if possible. Shardlow et al. [39] is perhaps the closest to our approach in that it attempts to link an author’s findings to the author’s statements of intended knowledge gain, although that is a more limited set of knowledge goals than we pursue.

As described in more detail below, we adopt the lexical cues used in the hedging literature and extend them to encompass a broader collection of ignorance phenomena. Note that hedging, while a related concept, is not the same as an ignorance statement: “The exact molecular function of SEPW1 protein is unknown to date” is an ignorance statement, but is not hedged, while “Thus, depending on the cellular environment, the short- and long-term effects of Tax expression can be quite different” is hedged [42], but not an ignorance statement.

## 2 Methods

To achieve our goal of identifying and characterizing questions stated in the biomedical literature, we must first formalize what it means to be an ignorance statement and the knowledge goal it entails, and then demonstrate that such ignorance statements exist and can be identified, either manually or automatically, in the literature. To do this, we 1) develop a taxonomy of ignorance statements, 2) create annotation guidelines for recognizing such statements, 3) validate their use through an annotation task to create a manually labelled corpus, and lastly, 4) use the corpus to benchmark classifers to automatically recognize ignorance statements (see Figure 1. These resources help ground future research aimed at exploring the state of our scientific ignorance.

**Fig. 1:**
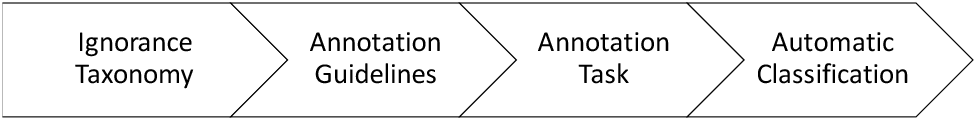
Methods Flowchart. A flowchart of the methods in navy with the steps in gray.

### 2.1 Materials

Scientific articles from our subject area of prenatal nutrition were taken from The PubMed Central Open Access (PMCOA) subset of PubMed [1], to ensure access to full-text articles and free sharing of data. We queried PMCOA using 54 regular expressions determined in consultation with a prenatal nutrition expert, which included keywords such as {prenatal, perinatal, and antenatal} paired with keywords like {nutrition, vitamin, and supplement} (the full query can be found on the GitHub page below). In total we gathered 1,643 articles, subsets of which were used for each task below. All computation was written in Python 3, with its associated packages. In addition, the annotation task used Knowtator [36] and Protege [22]. All code and associated materials, including the full query, taxonomy and annotation guidelines can be found here: https://github.com/UCDenver-ccp/Ignorance-Question-Work.

### 2.2 Ignorance Taxonomy

To better understand how to identify and categorize ignorance statements, we first manually reviewed a subset of 736 article abstracts among the 1,643 prenatal nutrition articles from PMCOA. Specific words or phrases, known as **lexical cues**, in each sentence were highlighted that signified a statement that knowledge was incomplete or missing (Figure 2). These lexical cues were then grouped together and organized into a taxonomy of labeled ignorance statement categories based on the knowledge goal each statement suggested. Further lexical cues and categories were inspired by and added from existing work [14, 23, 34, 16, 41, 20, 21, 42] to create an initial **ignorance taxonomy** driven by knowledge goals. The majority of lexical cues correspond to a single ignorance category, and thus imply a specific category assignment, though some cues, such as *challenge, if so*, and *imply* appear in multiple categories and the correct category assignment then depends on sentence context. The taxonomy contains a hierarchy of broad and narrow categories based on the different types of knowledge goals. Lastly, the taxonomy is dynamic and iteratively updated further during the annotation task, described below.

**Fig. 2:**
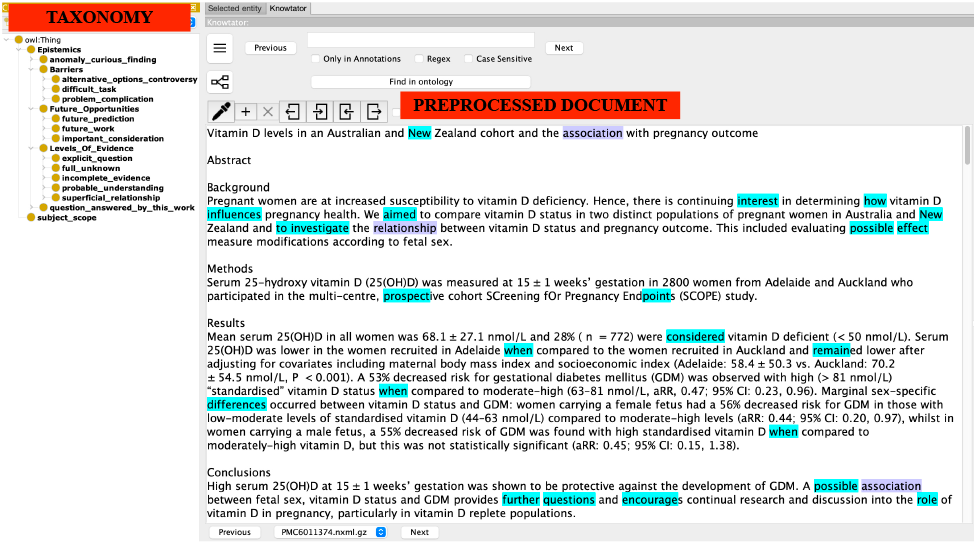
Example of Initial Document Markup

### 2.3 Annotation Guidelines

A set of annotation guidelines were developed that formalize the process that emerged during the manual review of abstracts to build the taxonomy to determine if identifying ignorance statements is reproducible and feasible. In particular, we recognized the importance of lexical cues and example sentences to help guide the annotators. Thus, the guidelines describe the annotation process beginning with text where lexical cues were automatically highlighted, in order to focus the annotator, rather than starting from unmarked full-text articles. The annotation task then becomes to determine whether the sentence containing a given highlighted lexical cue is indeed a statement of ignorance, to ensure that the ignorance category implied by the lexical cue is the correct one, and to identify additional ignorance statements and their lexical cues that are not yet part of the taxonomy (not highlighted already).

An important aspect of helping annotators reliably identify whether a lexical cue signifies an ignorance statement or not is through the use of examples. We gathered and reviewed 150 sentences from the general PMCOA to provide both positive and negative examples for each lexical cue identified. For example, “<however, there is <u>CONTRADICTORY</u> evidence from recent studies regarding the influence of IL-6 on insulin action and glucose metabolism> [4546]” represents a positive example of an *alternative option/controversy*, where we need to resolve disagreements about the influence of IL-6. At the same time, the sentence: “although yin and yang are <u>CONTRADICTORY</u> in nature, they depend on each other for existence” is a negative example, explaining the relationship between yin and yang. The annotation guidelines thus contain the ignorance taxonomy, along with all definitions and lexical cues for each taxonomy category, and include both positive and negative examples of ignorance statements. The annotators reference these examples throughout the annotation task.

In order to capture an ignorance statement completely, not only do we want the lexical cue and taxonomy category, but we also want the **subject** of it: the biomedical knowledge the lexical cue is qualifying. For that, we borrowed and modified the scope guidelines from BioScope [42]. Our annotations are different, but the scoping principles are similar: to capture the sentence fragment that is the subject of the task at hand. We chose to capture the full sentence that contains a lexical cue as the subject due to difficulties in capturing just fragments. Identifying these subjects will then allow for future work to identify the specific biomedical concepts that these ignorance statements refer to using existing automatic concept recognition tools (e.g., [4]). In the example above, the brackets < > signify the subject of the ignorance statement, i.e. the full sentence. Note that there are no brackets around the negative example as it is not an ignorance statement. The annotation guidelines describe how to remove the annotation for the incorrectly highlighted cues and how to add the subject by highlighting the full sentence for the true ignorance statements. For the full guidelines see: https://github.com/UCDenver-ccp/Ignorance-Question-Work.

### 2.4 Annotation Task

The goal of the annotation task was to produce a high quality, manually labelled dataset of full text documents. In brief, all documents were preprocessed to highlight all lexical cues, as well the ignorance category implied by the cue according to the taxonomy (or the first category if multiple apply). Then two independent annotators, Mayla R. Boguslav (MRB) and Elizabeth K. White (EKW) annotated 4-7 documents at a time by identifying all ignorance statements, indicating their ignorance category and subject, and then evaluated the quality and agreement of their annotations per batch using inter annotator agreement (IAA) measures.

All disagreements were adjudicated to finalize the annotations for the gold standard corpus, adjusting the guidelines and taxonomy accordingly.

Knowtator [36] was used for the annotation process, where the lexical cues were pre-highlighted and the ignorance taxonomy was built into Protege [22] like an ontology to be to browse all the definitions and lexical cue examples (Figure 2). For each pre-highlighted cue, each annotator independently decided whether it signified an ignorance statement. If not, then the annotator deleted the highlighted cue. If so, then the annotator left the cue highlighted, ensured the implied ignorance taxonomy category was correct (choosing the correct one if multiples), and highlighted the subject of the ignorance statement (i.e. the full sentence) (Figure 3). Notice the difference between the highlighted cues in Figure 2 and the gray highlights in Figure 3, in that some pre-highlighted cues were deleted and some were added. At the same time, the annotators read the full article and added new cues not already captured in the ignorance taxonomy, which were added to the taxonomy in the next round.

**Fig. 3:**
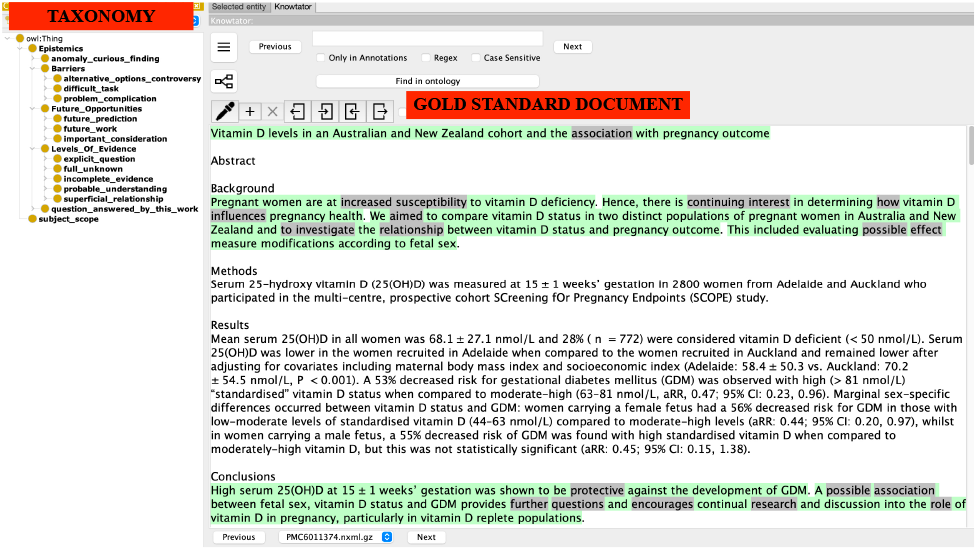
Example of Final Gold Standard Corpus Markup

To determine if the annotations are reliable, inter-annotator agreement (IAA) [18] was calculated once both annotators were done with their independent annotations, and the adjudication process began. The IAA is a measure of how well the annotations agree. Here, we calculated the F1 score between the two annotations, taking one annotation set as the “reference” and calculating precision and recall with the other one (note that changing the “reference” flips precision and recall, and thus F1 score would remain the same). The F1 score then was the harmonic mean between precision and recall. The IAA was calculated on the exact span of lexical cues or subjects chosen, as well as ignorance category assignments. We also calculate fuzzy IAA when the category assignments match but not the span of the cue or the subject, or vice versa.

Spans of the lexical cues can overlap when annotators recognized differing numbers of words for multi-word cues. For example, the annotators might agree on the ignorance category but highlight either <u>need</u> or <u>need to be</u> in the sentence “Thus doses of D vitamin and calcium supplementation, which may differ from those recommended in normal pregnancy, <u>NEED TO BE</u> carefully tailored in thyroidectomised patients.” We take the maximum span as the final between the two annotators (<u>need to be</u>). This can also occur with the subject annotations, where use of the Knowtator software may result in different spans highlighted.

Category assignments can overlap when one person annotated to the category implied by the specific lexical cue while the other annotated to a broader category. In the end we choose the narrowest applicable category in the taxonomy, to ensure we are capturing the correct information. All of these fuzzy matches are resolved during the adjudication process. Ideally, IAA should be over 80% to trust the annotations and reliability of the guidelines [37].

The final product of the annotation task is the gold standard corpus (see Figure 3). To finalize the annotations, the two annotators discussed all disagreements and these discussions led to updates in all or some of the the article annotations, the annotation guidelines, the ignorance taxonomy, and the lexical cue list. This process was repeated each time incorporating new updates.

### 2.5 Automatic Classification

To determine if the annotation task is feasible to automate, we test standard classifiers, with default settings, using the manually-labelled dataset to evaluate performance(see Figure 5. Our purpose is to demonstrate feasibility, not develop the optimal model, so a larger investigation of algorithms and tuned parameters will be the subject of future work with a larger corpus. Classification can be made at the sentence or word level. At the sentence level, we first have the binary task to determine whether or not a sentence is an ignorance statement. We only captured each sentence once as long as it had at least one lexical cue, even though one sentence can contain multiple. If so, we then have the multi-classification problem to determine the specific taxonomy category, or multiple categories if the sentence contains multiple lexical cues, as in Figure 4. For each predicted ignorance statement, we can also classify whether each word of the sentence is part of a lexical cue or not, and if so, for which ignorance category. For all classification tasks, we used a 90:10 split for the training and testing data and evaluated the models using F1 score.

**Fig. 4:** Annotation Task Example in Knowtator. An example document as first presented to the annotators, preprocessed to highlight the putative lexical cues (Figure 2) and then the same document as it appears in the finalized, abjudicated gold standard set, with the green highlight indicating the subject and the gray highlights indicating the true ignorance cues (Figure 3).

**Fig. 5:**
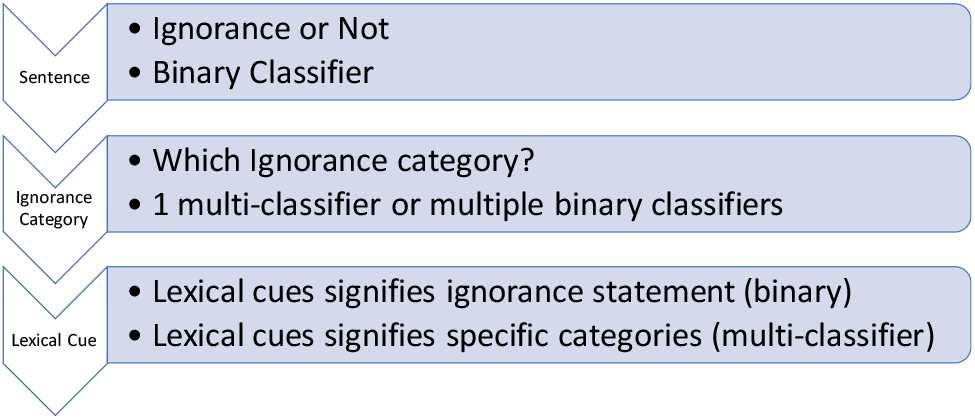
Classification Flowchart. A flowchart of the different classification problems.

For the binary sentence classification, we initially compared three different algorithms: logistic regression (LR), Gaussian Naive Bayes (GNB), and artifical neural networks (ANN) [35, 11]. Results from LR and GNB models were highly comparable to using ANN (data not shown) which we adopt here, due to the flexibility of ANN to build non-linear decision models [12]. Full-text documents from PMCOA were first split into sentences, which were then processed by the CountVectorizer package in python to tokenize words, remove punctuation and special characters, and produce a count vector of words as input to the ANN. The final tuned ANN had only 2 dense layers with dimensions 50 (to be proportionate with the large input shape) and 1. Overfitting was avoided by adding the training truncating functions (epochs ranged between 6 to 12). The batch size of 16 chosen for the training was small to allow for faster training and better generalization [10]. We also applied the stratify function to make sure that the data splitting for the test is equal. Due to an imbalance of our data, we balanced it to the class with fewer samples before training to achieve a stable learning process and verified this decision by training on the non-balanced data in a separate trial.

The sentence multi-classification problem, classifying each ignorance sentence into a taxonomy category, has a similar setup to the sentence-binary task, using a vector of word counts as inputs. This task though is more complex because some sentences map to multiple categories, there can be overlapping lexical cues, the categories are not balanced, and multi-classification problems are inherently harder than binary tasks. To combat all of these problems, we decided to create a binary classifier for each taxonomy category that classifies all sentences as either the category of interest or not (meaning the other taxonomy categories). All classifiers, one for each taxonomy category, are run over all the data, allowing for one sentence to be classified in multiple categories easily.

The word-level classification tasks are to determine which words in a sentence indicate a lexical cue (binary task), and specifically for which taxonomy category (multi-classification). As our taxonomy is very similar to an ontology, we can use prior methods for concept recognition [4] that explores different algorithms over many different ontologies. In particular, we make use of the best performing span detection algorithms, namely CRF [24] and BioBERT [26], to determine the words in all lexical cues given an article. The CRF and BIOBert are trained for each specific iteration of the word-level classification. Thus, we first word-tokenize the articles using WordPuctTokenize in Python. We then assign each word a BIO-tag based on whether the word is at the beginning of a lexical cue (B), inside of it if it is a multi-word cue (I), outside of it meaning not a word in the lexical cue (O), or if the lexical cue contains a discontinuity (e.g., *no*…*exist* where the “…” signifies a discontinuity), we label the words that exist between the lexical cues (O-). The input to the CRF and BioBERT to train then are the words with their target BIO-tag label. When predicting, the input are the word-tokenized articles and the output is a BIO-tag for each word. To then determine what the lexical cue is, we re-assemble the BIO-tags by finding the B tags (single word lexical cues), combining the words with B and I in a row (multi-word lexical cues), ignoring the O labels (not lexical cues), and by combining B, I, and O- (discontinuous lexical cues). A more thorough discussion of BIO-tags can be found in [4].

For the word-level binary task, we take every lexical cue from all ignorance categories, classify the words into BIO-tags, and then re-assemble them into the lexical cues. Since we are on the word-level, the binary classifier can find new lexical cues that are not already in our lexical cue list by finding new combinations of words for example. So, we can compare the predictions of the classifier to our set of lexical cues in the taxonomy to determine if the classifier is finding new ones.

For the word-level multi-classification task, we explore two different methods. First, we created a model (CRF and BIOBert) for each ignorance taxonomy category following the previous work [4] because of the same issues stated above for the sentence multi-classification problem. At the same time, however, BIO-tags have the capability to encode which category the word is from (i.e., B-*taxonomy category* or I-*taxonomy category*), and so we can create one multi-classifier (CRF or BIOBert) for the word-level classification as well.

## 3 Results

### 3.1 Statements of ignorance employ a rich vocabulary

Manual review of 736 paper abstracts to develop a taxonomy of ignorance statements and their implied knowledge goals revealed that abstracts contained on average seven (minimum 0, maximum 24) ignorance statements, involving 900 lexical cues. Subsequent refinement of 60 full text articles during the annotation task contributed another thousand lexical cues or so. Organizing the lexical cues by their knowledge goals lead to the final taxonomy with five broad categories, composed of 13 narrow ones:

- **Question Answered by this Work**
- **Levels of Evidence:** full unknown, explicit question, incomplete evidence, superficial relationship, probable understanding
- **Anomaly/Curious Finding**
- **Barriers:** alternative options/controversy, difficult task, problem/ complication
- **Future Opportunities:** future work, future prediction, important consideration

Note that both *question answered by this work* and *anomaly/curious finding* are considered as categories at both the broad and narrow level. As for the three broad categories that unify multiple narrow categories, “Levels of Evidence” contains statements that answer the questions of how much evidence we have and how confident we are in that evidence, in increasing order. “Barriers” contains statements of obstacles, complications, or multiple options that prevents research from moving forward and needs to be overcome. “Future Opportunities” includes statements of future needs such as future work or considerations. These broad categories help simplify the 13 categories for annotation purposes. For the 13 narrow categories, we collected 1,890 lexical cues that can signify an ignorance statement after the full annotation task (Table 1). This signifies that the language used to articulate similar knowledge goals is rich and varied. *Incomplete evidence* contains the most cues. Interestingly, most ignorance statements are not presented as an explicit question with a question mark or a question word (who, what, where, etc.), yet still imply a question and still present a knowledge goal as *explicit question* has the fewest number of cues. The remaining 12 ignorance categories are driven by their knowledge goals, from finding more evidence for *incomplete evidence* to creating new methods for a *difficult task*.

**Table 1.** The ignorance taxonomy with definitions, knowledge goals, and example cues. The categories in bold are both broad and narrow categories.

| Category | Definition | Knowledge Goal | Example Cues | Total Cues |
| --- | --- | --- | --- | --- |
| **question answered by this work** | A statement of a goal or objective of a study that is attempted or completed during the study. | to find the answer(s) in the article; determine if the question(s) is (are) fully answered in the article | aim, goal, objective, our study, sought, to determine | 58 |
| full unknown | A statement that indicates something is not known (a lack of information), or information is presented for the first time (new or novel) and a significant amount of research is needed; not a statement about the absence of something. | to explore the unknown further to gain any insights | could not find, don't know, elusive, not...established, uncertain, still unclear | 137 |
| explicit question | An explicit statement of inquiry (with a question mark or question word such as how, where, what, why). | to find answers to the question and/or discover methodologies that will help answer the question | ?, what, where, wondered, why | 17 |
| incomplete evidence | A positive or negative statement proposing a possible/feasible explanation for a phenomenon on the basis of limited evidence as a starting point for further investigation OR a statement that information is needed to support an assertion or claim, including both positive and negative statements. Either a statement that some evidence already exists, explaining how current findings support previous work, adding confidence to a claim OR a statement that information is limited, more research is needed or is ongoing including limitations – biases or short comings related to the study design and execution. | to gather more evidence to support the claim OR conduct more research to determine the validity of the claim; complete the partial picture; consider the short comings and biases for the next experiment and how it can be addressed. | a good understanding, believe, evidence...limited, has been suggested, hypothesis, no studies, possibly, preliminary stage, remains under investigation, still being discovered, support, trend | 619 |
| superficial relationship | A statement about a connection, link, or association between at least 2 variables; connectedness between entities and/or interactions representing their relatedness or influence. | to confirm the connection, link, or association between variables; determine the full underlying relationship between variables | affect, associated, correlate, factor, influence, interact, link, pattern, tend | 133 |
| probable understanding | A statement staking a claim to the most likely explanation, relationship, or phenomenon; assumes that there is a good chance this understanding is correct. | to determine if the most likely option is correct or if it's another option is more feasible | almost all, assumed, concluding, evident, it is clear, most likely, thus | 119 |
| **anomaly/curious finding** | A statement of a surprising result, conclusion, observation or situation; the researchers were not expecting the result, conclusion, observation or situation but are intrigued by it. | to explore the surprising result, conclusion, or situation more and determine if the result, conclusion, observation, or situation are repeatable | appeared to be, interestingly, noteworthy, surprisingly | 89 |
| alternative options/controversy | Either an explicit statement of multiple (at least 2) choices, actions, approaches, or methods that need to be experimentally determined, including statements with an implied second option, such as “whether”. This includes a statement of disagreement amongst researchers OR a lack of consensus OR at least two possible answers presented as results from different researchers - usually in reference to previous results and stated when results disagree with each other OR contradictions. | to determine the correct option or a better option and if there are disagreements, or to determine the truth to break any disagreements | cannot rule out, claims, has been challenged, whether, whilst | 193 |
| difficult task | A statement of something not easily done, accomplished, comprehended, or solved; or a complicated thing with a multitude of underlying pieces or parts; heterogeneity; excludes medical complications. | to create methods to study the complicated system and to better understand any piece of the complicated system; potentially requires new experiments or better techniques | not feasible, remains...challenge, variability, rarely able to | 69 |
| problem/complication | A statement of issues, problems, mistakes, or medical complications that are cause for anxiety and/or worry. | to determine the gravity of the concern and determine if it needs to be dealt with before the next experiment or study | issue, error, insufficient, lack of reproducibility, publication bias, underestimated | 86 |
| future work | A statement of extensions, including next steps, directions, opportunities, approaches, or considerations of the described work that may be implemented at some future time point. This also includes a statement of suggestion or a proposal as to the next best course of action, especially one put forward by an authoritative body; advice telling someone the best action to take. | to determine the next course of action based on this future work proposal | additional research, are needed, continue to explore, further study, more...studies, recommend, warrants, worthy of closer attention | 201 |
| future prediction | A statement of extrapolation of given data into the future and/or from past observations, without reference to next steps. | to run the simulation or experiment to determine if the prediction is correct; publicize the outcomes of the study to the correct people | allow, expect, if so, serve as a basis, will | 17 |
| important consideration | A statement calling for attention including an action needed to be taken immediately or information that needs to be disseminated immediately OR critical: being in or verging on a state of crisis or emergency OR urgently needed OR absolutely necessary. | to take the urgent action ASAP or distribute the knowledge ASAP | call for action, cautious, crucial, emphasis, global problem, high on the agenda, necessary, relevant to note, vital | 152 |

### 3.2 Robust annotation guidelines yield a high quality corpus

Annotators were asked to identify ignorance statements in 60 full-text articles about prenatal nutrition from PMCOA in batches of 4-7 articles for a total of 13 batches. Annotation guidelines were provided that specified the ignorance taxonomy, all definitions and lexical cues for each taxonomy category, and included positive and negative examples of ignorance statements for each category.

The positive and negative examples for each lexical cue were very helpful in making the annotation task more feasible. Surprisingly, almost every cue contained a negative example. Even for seemingly obvious cues such as <u>unknown</u>, there is a negative example. First, a positive example is “*<*it is <u>UNKNOWN</u> what triggers the bidirectional transcription of sine b2, but boundary activity only occurred when both transcription machineries were in play*>*.” But a statement such as “to make inference, the maximum likelihood method is applied to estimate the <u>UNKNOWN</u> parameters in the empirical log-odds ratio models given in (3)-(6)” is a negative example in that it is a description of a methodology that helps determine missing parameters. The guidelines were updated during the annotation task; as new cues were identified they were added to the guidelines and the statements they were found in were used as positive examples. Thus, some cues only have positive examples and the guidelines are not completely static.

After each of 13 batches, inter-annotator agreement (IAA) was assessed for concordance in span indicated for the lexical cue or subject and for category assignment, calculating both exact match IAA and fuzzy match IAA for each (Figure 8. For the first six batches, the category IAA was higher than the subject, and then it switched, probably due to the switch from annotating sentence fragments to full sentences as the subject. Also, over time, the annotators were more likely to recognize the same sentences as ignorance, even if different lexial cues were chosen. Thus by batch six, both the category and subject IAAs plateaued until the last batch. The last batch IAA dropped a small amount, likely because it contained three articles that differed from the rest in style and topic. Overall, we achieved IAA around 80% for the ignorance category and 90% for the subject (Figure 6, indicating that the annotation guidelines were convergent and robust.

**Fig. 6:**
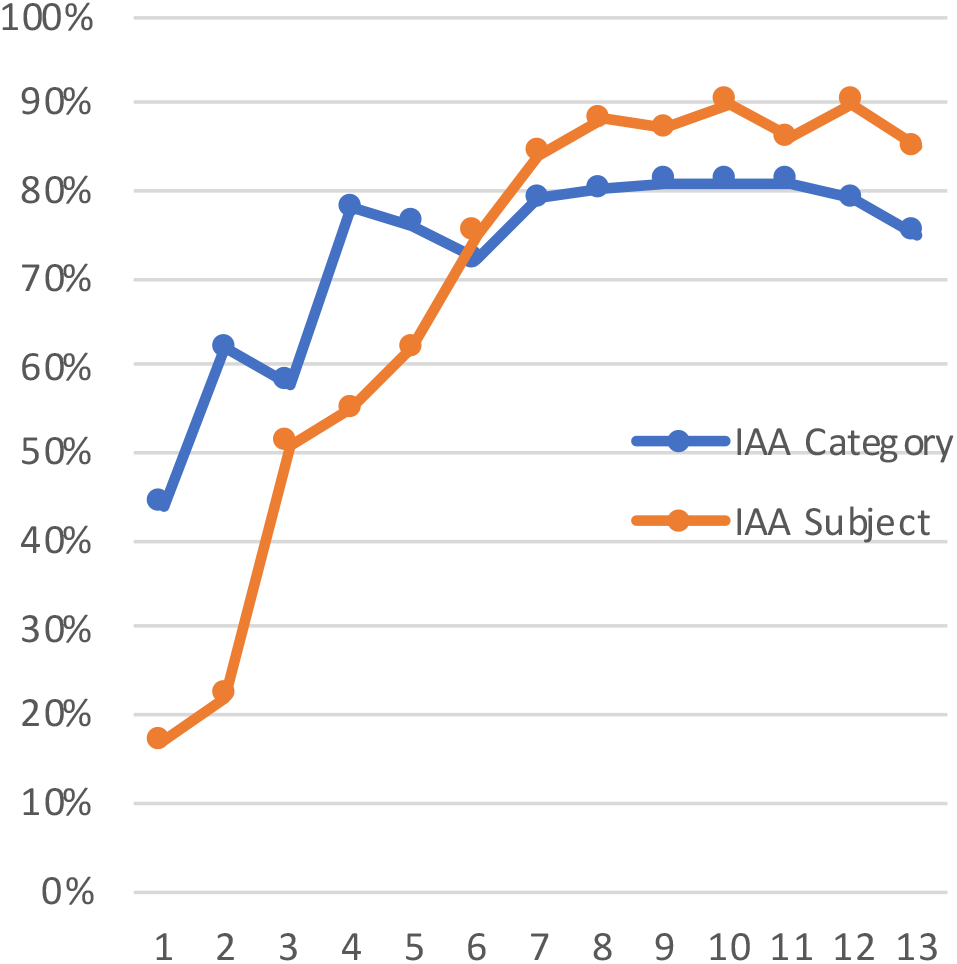
Exact Match

**Fig. 7:**
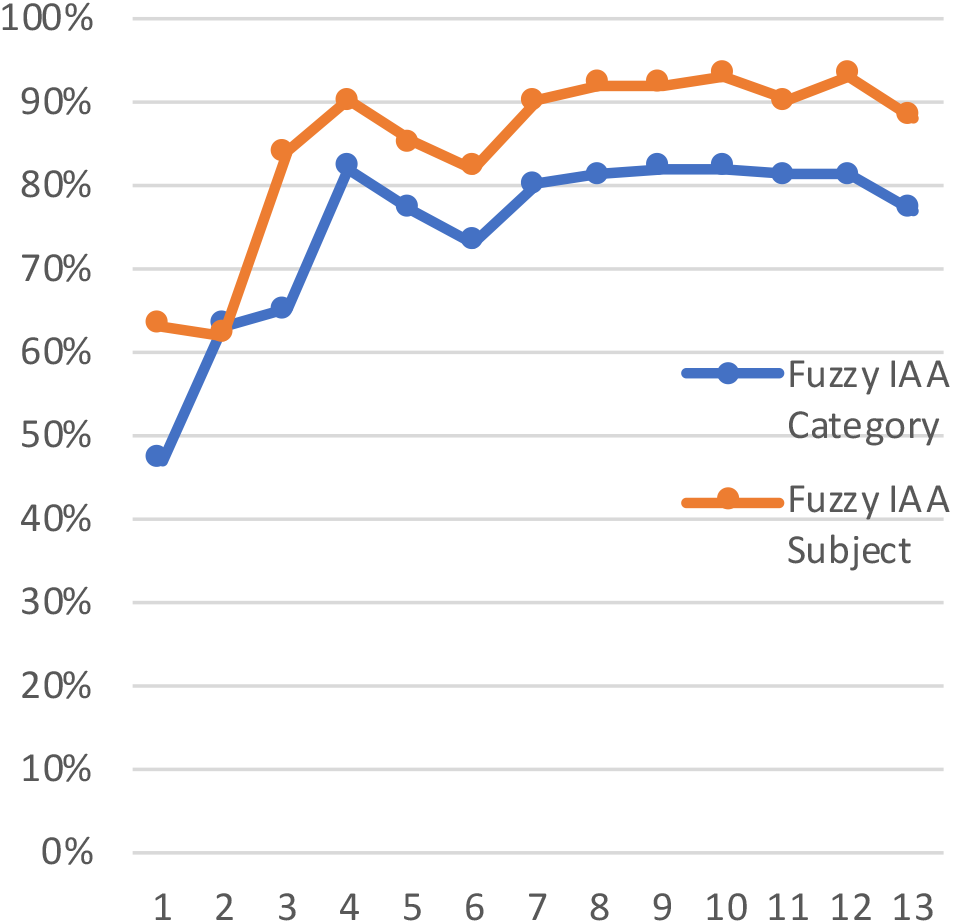
Fuzzy Match

**Fig. 8:** IAA statistics for Ignorance Annotations by Batch. Plot of IAA vs. of article batches (approximately bimonthly) for annotation of the corpus with both ignorance category (blue) and subject (orange) annotations. IAA has been calculated as F1-score, requiring (a) exact or (b) fuzzy matches.

Allowing for both kinds of fuzzy matches greatly increases the IAA for subject annotations, especially for the first six batches, smoothing out the curve (see Figure 7. The overlapping match specifically boosted it here. The subject IAA is now generally higher than the category IAA except for the second batch. The category IAA increases slightly but maintains its curve and now aligns well with the subject IAA curve. Thus, the fuzzy match greatly improves subject IAA, slightly improves category IAA, and better aligns the curves to each other.

Overall, we present a high quality corpus with robust annotation guidelines. Combining all articles, we can reliably annotate ignorance statements: we achieve a category IAA of 78%, subject IAA of 87%, fuzzy category IAA of 79%, and fuzzy subject IAA of 90%.

### 3.3 The scientific literature is rich in ignorance statements

The scientific literature, specifically in prenatal nutrition, is rife with ignorance statements (see Table 2). There are over 10,000 lexical cue annotations included in nearly 4,000 subject annotations in 60 articles. These articles include 7,304 sentences and 249,133 words. The majority of these category annotations refer to *incomplete evidence* or *superficial relationship*. For each ignorance category, the average number of annotations per article ranges from 1 to 60, with the median from 1 to 40. Also, not every article includes annotations to every ignorance category (the minimum number of annotations per article is zero). However, there are articles with over 100 annotations to one category. Thus, each article contains many ignorance statements that map to all the taxonomy categories.

**Table 2.**
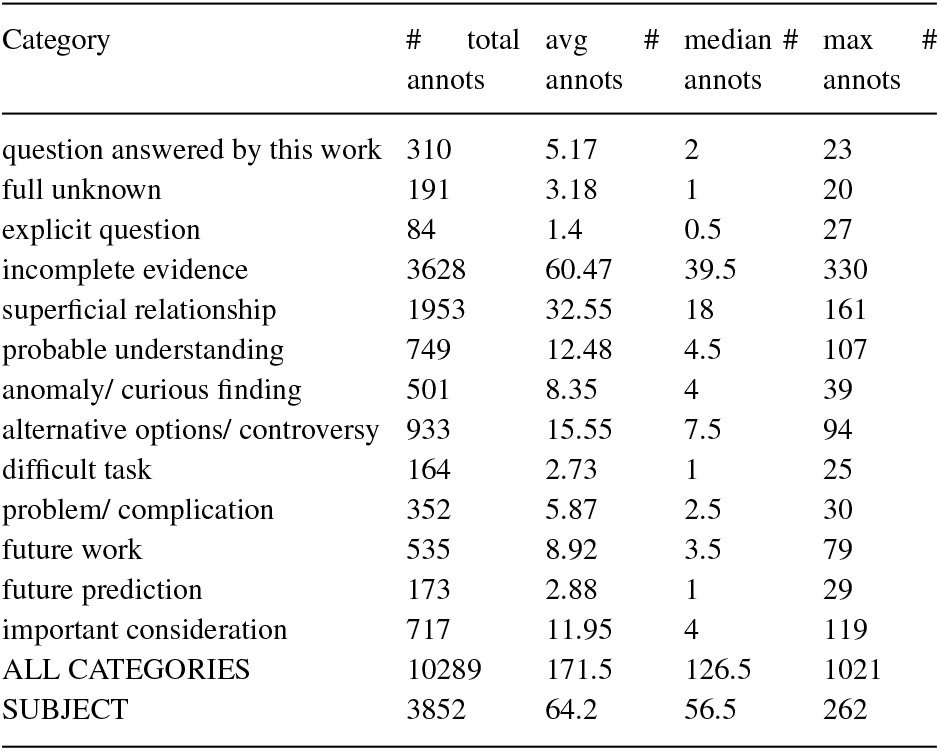
Per Article Counts of Annotations. Total number of annotations in all articles and statistics per article. Note that all categories except for ALL_CATEGORIES and SUBJECT (have 1) have zero minimum number of annotations. Note annots = annotation, avg = average, and max = maximum.

Even with so many lexical cue annotations, many of them are repeats of the same cue (see Table 3). Only 8% of the 10,000 annotations are unique lexical cues. Further, only 43% of all cues collected (nearly 2,000) are used in the articles. Still, the trends of the unique counts are the same as the full counts of annotations. *Incomplete evidence* and *superficial relationship* still comprise the majority of the annotations. There remain articles without annotations to every ignorance category. Lastly, there are articles with over 40 unique annotations to one category. Thus, even looking at unique annotations, each article contains many unique lexical cues to all categories.

**Table 3.**
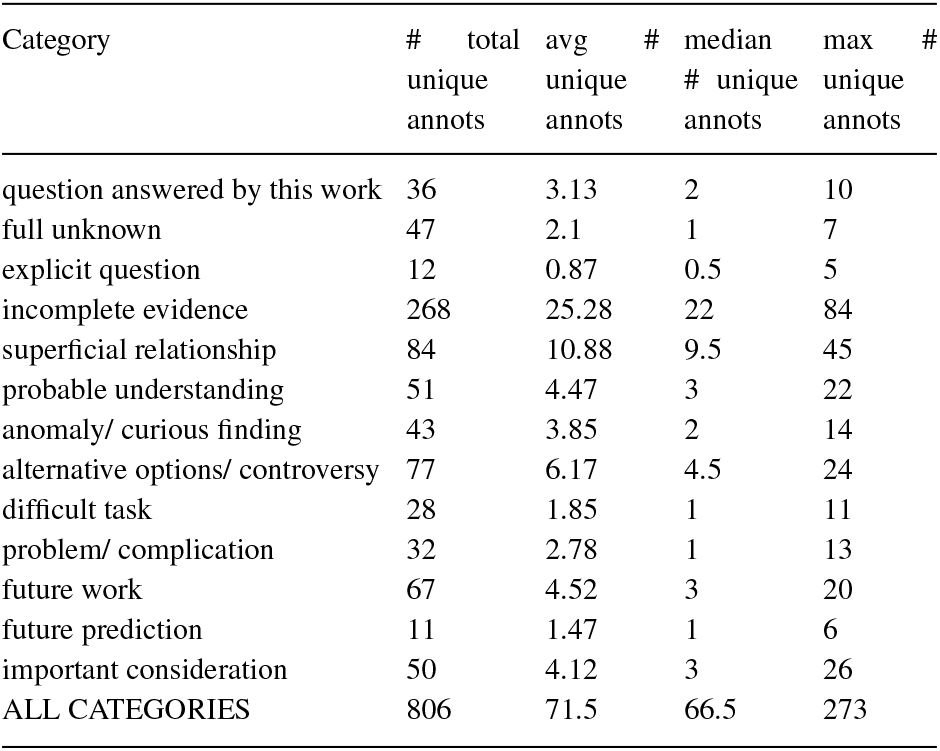
Per Article Unique Counts of Annotations. Total number of unique annotations in all articles and statistics per article. Note that all categories except for ALL_CATEGORIES (has 1) have zero minimum number of unique annotations. Note annots = annotation, avg = average, and max = maximum.

The frequency of lexical cue annotations varies based on article section (see Table 4). Unsurprisingly, the conclusion contains the most annotations, followed by the discussion and then the method section. The abstract section contains the fewest annotations, which may be due to the normally small size of this section compared to the others. On average, each section has at least 2 annotations and at most 117 annotations. The medians are slightly lower, indicating some outlier articles with many annotations. In fact, the maximum number of annotations in one section in one article is 524 in the conclusion section. Aside from the method and conclusion sections, there exist articles with annotations in some sections but not all. For each article with a conclusion section, there are at least four annotations. Similarly, every article with a method section has an annotation in that section. Thus, looking within the article, the different sections contain differing numbers of ignorance statements.

**Table 4.** Per Article Annotation Counts per Section. Total number of annotations by section in all articles with section delineation and statistics per article. Note annots = annotation, avg = average, min = minimum, and max = maximum.

| Section | # articles | # total annots | avg | # median | # min | # max |
| --- | --- | --- | --- | --- | --- | --- |
| abstract | 42 | 80 | 1.9 | 1 | 0 | 29 |
| introduction | 55 | 1416 | 25.75 | 14 | 0 | 571 |
| methods | 35 | 1403 | 40.09 | 30 | 1 | 367 |
| results | 29 | 323 | 11.14 | 6 | 0 | 46 |
| discussion | 31 | 1940 | 62.58 | 37 | 3 | 258 |
| conclusion | 28 | 5127 | 183.11 | 107.5 | 4 | 990 |

### 3.4 Ignorance statements and lexical cues can be automatically identified

We can automatically identify ignorance statements (see Table 5) and lexical cues (see Table 6. Due to limited space however, we only show the top algorithms for classification with both the training and testing F1 scores. For binary sentence classification (is a sentence an ignorance statement or not no matter the ignorance category), we achieved an F1 score of 0.85 with 7 epochs for the ANN.

**Table 5.**
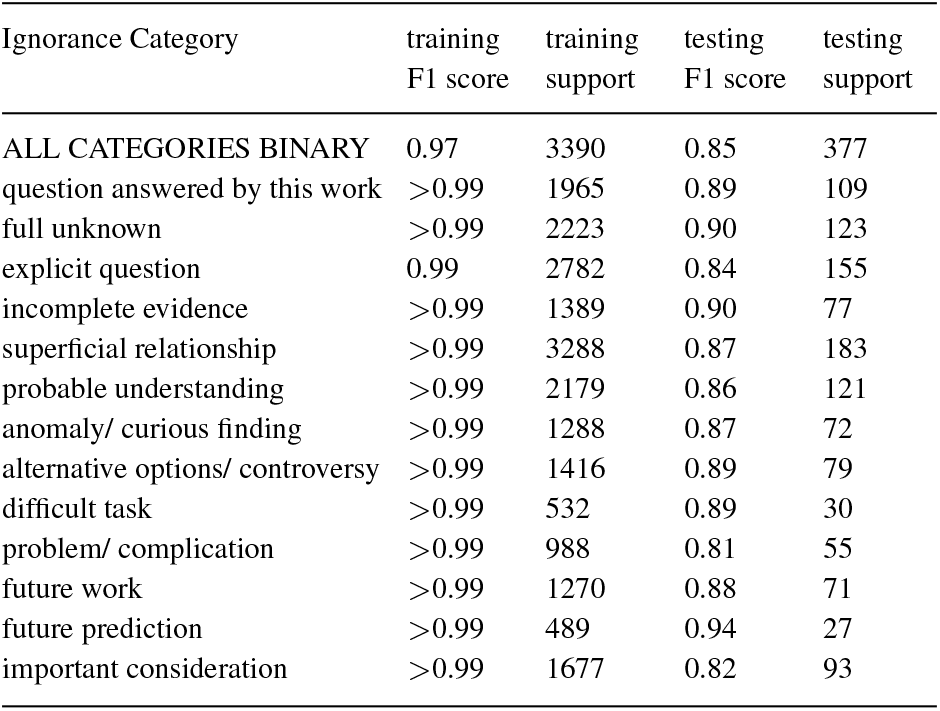
Sentence Classification both binary and all 13 categories. Note that one sentence can map to more than one category and so they will not add up to the total binary.

**Table 6.** Word Classification both all together, binary and to all 13 categories.

| Ignorance Category | training F1 score | training support | testing F1 score | testing support |
| --- | --- | --- | --- | --- |
| ALL CATEGORIES BINARY | 0.95 | 11552 | 0.93 | 1210 |
| question answered by this work | 0.91 | 474 | 0.83 | 51 |
| full unknown | 0.85 | 299 | 0.8 | 28 |
| explicit question | 0.98 | 68 | 0.9 | 15 |
| incomplete evidence | 0.96 | 4342 | 0.96 | 520 |
| superficial relationship | 0.98 | 1812 | 0.99 | 199 |
| probable understanding | 0.96 | 753 | 0.95 | 73 |
| anomaly/ curious finding | 0.91 | 522 | 0.94 | 61 |
| alternative options/ controversy | 0.92 | 961 | 0.9 | 117 |
| difficult task | 0.87 | 199 | >0.99 | 19 |
| problem/ complication | 0.93 | 415 | 0.94 | 36 |
| future work | 0.91 | 716 | 0.87 | 84 |
| future prediction | 0.97 | 180 | 0.94 | 18 |
| important consideration | 0.98 | 737 | 0.97 | 85 |
| ALL CATEGORIES COMBINED | 0.84 | 9416 | 0.85 | 1073 |

For the sentence multi-classification problem, we created 13 binary models with the target class against the other 12 classes (e.g., *difficult task* class vs. the rest of the classes). As each category had fewer positive examples than the other categories combined, we balanced the data and noticed that in the non-balanced data the F1 score is biased towards the class that has the higher number of samples. For the balanced trials, testing F1 scores ranged between 0.8 and 0.94 (see Table 5).

Going deeper to identify the specific lexical cues using BIO-tags on the word level, BioBERT performed better than the CRF for all word-level classification tasks (see Table 6 reporting only BioBERT). The binary task (ALL CATEGORIES BINARY) achieved an F1 score of 0.93 and contains the most data of all the tasks. Further, in general, all these lexical cues mapped to our current list of lexical cues in the taxonomy, except for about 6%. For the 13 binary classifiers created for the multi-classification task to each narrow category, the F1 scores ranged between 0.8 to greater than 0.99 based on varying amounts of data per category. Combining all this data into one multi-classifier using the BIO-tag capability to add a category (e.g., B-*incomplete_evidence*), we achieved an F1 score of 0.85. More data is necessary to truly validate these results but they do seem promising to show that we can automatically identify statements of ignorance

For all tasks, F1 scores are above 0.8 and rise even higher for categories with more data. Further, both the combined and binary methods performed quite well. For normalizing the binary method over all the data including training and testing, all lexical cues predicted mapped to the taxonomy except for about 6%. As the dictionary of cues map each cue to only one ignorance category, this number is quite low and contributes to an F1 score of 0.90 to all categories. The errors seemed to be due to minor variations on the ignorance cues already in the taxonomy, such as plurals or words in different orders. More data is necessary to truly validate these results, but they do seem promising.

## 4 Discussion

Identifying and categorizing ignorance statements in terms of their knowledge goal is an important new NLP task that formalizes and extends prior work in uncertainty, hedging, speculation, and meta-knowledge. Our ignorance taxonomy subsumes prior work by Firestein [14], who elucidates a few types of ignorance informally including curiosity, possibility, and controversy, which extend to *anomaly/curious finding, incomplete evidence*, and *alternative options/controversy* in our taxonomy. Kuhn [23] discusses anomalies that drive the crises, which is *anomaly/curious finding* in our taxonomy. Pearl et al. [34] aimed to forge connections from correlation to causation, the *superficial relationship* category in our taxonomy, since the literature is filled with associations. Han et al. [16] focused on healthcare with categories including probability, ambiguity, and complexity extending to *probable understanding, anomaly/curious finding*, and *difficult task* in our taxonomy respectively. Smithson’s [41] taxonomy includes incompleteness and probability, which map to *incomplete evidence* and *probable understanding* in our taxonomy. Kilicoglu et al. [21] using SemRep with categories including probable and possible, which fall under *probable understanding* and *incomplete evidence* in our taxonomy. Many of these works also provide lexical cues to help do their tasks, which we extend into nearly 2,000 cues. To the best of our knowledge this is the largest lexical cue list and shows that ignorance statements employ a rich vocabulary.

Using our ignorance taxonomy and lexical cues, we provide robust annotation guidelines that yield a high quality corpus. Human readers can reliably identify and categorize ignorance statements as demonstrated in our annotation task (see Figures 6-7). Further, this corpus highlights how rich the scientific literature is with ignorance statements (see Tables 2-4), both by article and by section. Also, most ignorance statements are implicit (the *explicit question* category is quite small), which makes this task difficult. As in prior work (e.g., [42, 13]), we provide a very detailed ignorance taxonomy, extensive annotation guidelines, and our gold standard corpus.

We also can automate this task by creating classifiers to identify ignorance statements and lexical cues. Although we did not exhaustively train and tune our models, our results look promising that this task is feasible (see Tables 5 and 6). We achieved F1 scores above 0.8 for all classification tasks. Further, in the binary word-level classification task to identify any lexical cue that signifies an ignorance statement, we may be generating new lexical cues beyond our list, as 6% of the predicted cues did not match our list. Future work can both explore classification algorithms more and these possible new cues.

The major limitation of this work is the small amount of data. The corpus contains 60 articles which is enough to train classification models, but not enough to conduct an external validation. Future work currently underway includes more annotations to create a separate evaluation set. Even with this limitation though, our results are quite promising for both ignorance sentences and the specific lexical cues. It is both important to be able to identify sentences and cues to formally represent and compute over these sentences along with other biomedical ontology terms such as genes, proteins, and diseases. Further, if a sentence maps to more than one ignorance category, identifying the specific lexical cues can distinguish between the categories. All training and testing F1 scores are above 0.8, which according to James Pustejovsky et al. [37], is a perfect agreement level, showing that our data is reliable.

Even with these limitations, discussions of these ignorance statements can both refine and improve research questions, identify established facts, and facilitate the comparison of approaches for further research. These discussions of scientific questions are of interest to researchers, educators, publishers, and funders because they provide insights and directions for new research and may provide context for existing results. Formalizing and disseminating such ignorance statements in the scientific literature is an important new NLP task with the potential to greatly impact how we view the literature and scientific progress in general.

## 5 Conclusion

Focusing on scientific questions or ignorance statements in the scientific literature as a new NLP task will prove to be a significant and worthwhile endeavor in finding new avenues to gain insights and build new text mining tools. Here we not only showed that this task is feasible, but created an ignorance taxonomy, a gold standard corpus, and promising preliminary classification models. The goal is to help enhance literature awareness by creating an (UN)knowledge-base or ignorance-base (compared to a knowledge-base) of all the ignorance statements found in the literature for all to explore, including students, researchers, publishers, and funders.

## Acknowledgements

The authors would like to thank Harrison Pielke-Lombardo for his updates to the Knowtator tool, William A. Baumgartner Jr. for his help with the literature and taxonomy work, Teri L. Hernandez for her consultations on prenatal nutrition, Michael M. Bada for his guidance on annotation tasks and the taxonomy, and Katherine J. Sullivan and Tiffany J. Callahan for many discussions about this work.

